# The hippocampus generalizes across memories that share item and context information

**DOI:** 10.1101/049965

**Authors:** Laura A. Libby, J. Daniel Ragland, Charan Ranganath

## Abstract

Episodic memory is known to rely on the hippocampus, but how the hippocampus organizes different episodes to permit their subsequent retrieval remains controversial. According to one view, hippocampal coding differentiates between similar events to reduce interference, whereas an alternative view is that the hippocampus assigns similar representations to events that share item and context information. Here, we used multivariate analyses of activity patterns measured with functional magnetic resonance imaging (fMRI) to characterize how the hippocampus distinguishes between memories based on similarity of their item and/or context information. Hippocampal activity patterns discriminated between events that shared either item or context information, but generalized across events that shared similar item-context associations. The current findings provide novel evidence that, whereas the hippocampus can resist mnemonic interference by separating events that generalize along a single attribute dimension, overlapping hippocampal codes may support memory for events with overlapping item-context relations.

## INTRODUCTION

Memories for past events include information about who or what was encountered (“items”), as well as information about the place, time, and situation in which the event took place (“context”). For instance, seeing a friend’s dog (an item) at the neighborhood park (a context) might remind you of the time you played fetch with that dog in your friend’s backyard (same item, different context), or you might remember that, the last time you visited the park, you ran into a coworker (different item, same context). This example illustrates how the human brain can both distinguish between specific events and generalize across events according to similarity in item and/or context information (Johnson, Hashtroudi, & Lindsay, 1993; Mitchell & Johnson, 2009). Memory for item-context associations is known to rely on the hippocampus (Brown & Aggleton, 2001; Cohen, Poldrack, & Eichenbaum, 1997; Scoville & Milner, 1957; Vargha-Khadem et al., 1997), but how the hippocampus organizes memories to facilitate item and context retrieval is unclear.

One prominent model (Marr, 1971; Norman & O’Reilly, 2003; Rolls & Kesner, 2006; Yassa & Stark, 2011) holds that the hippocampus has a computational specialization for mapping different events onto orthogonal neural codes, a process known as pattern separation. Pattern separation is thought to prevent catastrophic interference between representations of events with similar features; when hippocampal pattern separation fails, episodic memory is predicted to break down (Norman, 2010; Norman & O’Reilly, 2003). The pattern separation account is consistent with results from single-unit recording studies in rats showing that spatial coding in the hippocampus dramatically “remaps” with slight changes in spatial context (Bostock, Muller, & Kubie, 1991; Guzowski, Knierim, & Moser, 2004; Leutgeb, Leutgeb, Moser, & Moser, 2007; Lever, Wills, Cacucci, Burgess, & O’Keefe, 2002; Wills, Lever, Cacucci, Burgess, & O’Keefe, 2005), but studies in rodents have not yet identified a role for the hippocampus in pattern separation of items independent of their spatial context. However, several recent human neuroimaging studies (Huffman & Stark, 2014; LaRocque et al., 2013; Liang, Wagner, & Preston, 2012) measuring the similarity of hippocampal voxel activity patterns across trials as an indirect measure of neuronal population coding (Kriegeskorte, Goebel, & Bandettini, 2006; Kriegeskorte, Mur, & Bandettini, 2008) have found that hippocampal coding tends to differentiate between similar items to a greater extent than cortical regions (Huffman & Stark, 2014; Liang et al., 2012). Further evidence (LaRocque et al., 2013) suggests that greater hippocampal differentiation between similar items during learning predicts greater subsequent memory for these items after a long delay. One limitation of the studies described above, however, is that context similarity was not also manipulated. Accordingly, it is unclear whether context affects pattern separation in the human hjppocampus.

A second framework for understanding hippocampal function suggests that, whereas item and context information are processed separately in parallel neocortical pathways (Ranganath & Ritchey, 2012; Ritchey, Libby, & Ranganath, 2015), the hippocampus specifically contributes to the representation of item-context associations (Davachi, 2006; Eacott & Gaffan, 2005; Knierim, Lee, & Hargreaves, 2006; Ranganath, 2010). Consistent with this idea, single-unit recording studies in rats have shown that hippocampal neuronal ensemble coding for items encountered in a particular spatial context is highly reliable across repeated exposures (Manns & Eichenbaum, 2009; McKenzie et al., 2014). Complementary results in humans have shown that hippocampal voxel patterns are similar when the same item is presented repeatedly in the same position within a temporal context (Hsieh, Gruber, Jenkins, & Ranganath, 2014; Libby, Hannula, & Ranganath, 2014; Ritchey, Montchal, Yonelinas, & Ranganath, 2015).

The “pattern separation” (Marr, 1971; Norman & O’Reilly, 2003; Rolls & Kesner, 2006; Yassa & Stark, 2011) and “items-in-context” (Davachi, 2006; Eacott & Gaffan, 2005; Knierim et al., 2006; Ranganath, 2010) hypotheses are complementary, in that the former addresses how the hippocampus organizes distinct events with respect to each other (i.e., pattern separation) and the latter focuses on the relative importance of item-context associations in episodic memory. However, a fundamental and currently unresolved question is how the hippocampus represents events that involve similar item and context information. A strong version of the pattern separation hypothesis would suggest that the hippocampus sharply differentiates between events, even if they share item or context information (e.g., different encounters with a dog in the park). A strong version of the items-in-context perspective, in turn, would suggest that the hippocampus assigns overlapping neural representations to events in which similar items were encountered in similar contexts.

To test these predictions, we used functional magnetic resonance imaging (fMRI) to examine hippocampal activity patterns during a memory test that required retrieval of both item and context information. Critically, we manipulated the similarity of both item and/or context information: Different exemplars from the same item category were learned in each of four different context locations, resulting in sets of events with overlapping item attributes, context associations, or both (Figure 1a). Participants were then scanned during a recognition test that required recall of item-context relations (Figure 1b). We analyzed the similarity of hippocampal voxel activity patterns evoked during successful retrieval of item-context relations to estimate the extent to which brain areas generalized or differentiated across memories with overlapping item, context, and item-in-context information (Figure 2).

**Figure 1.**
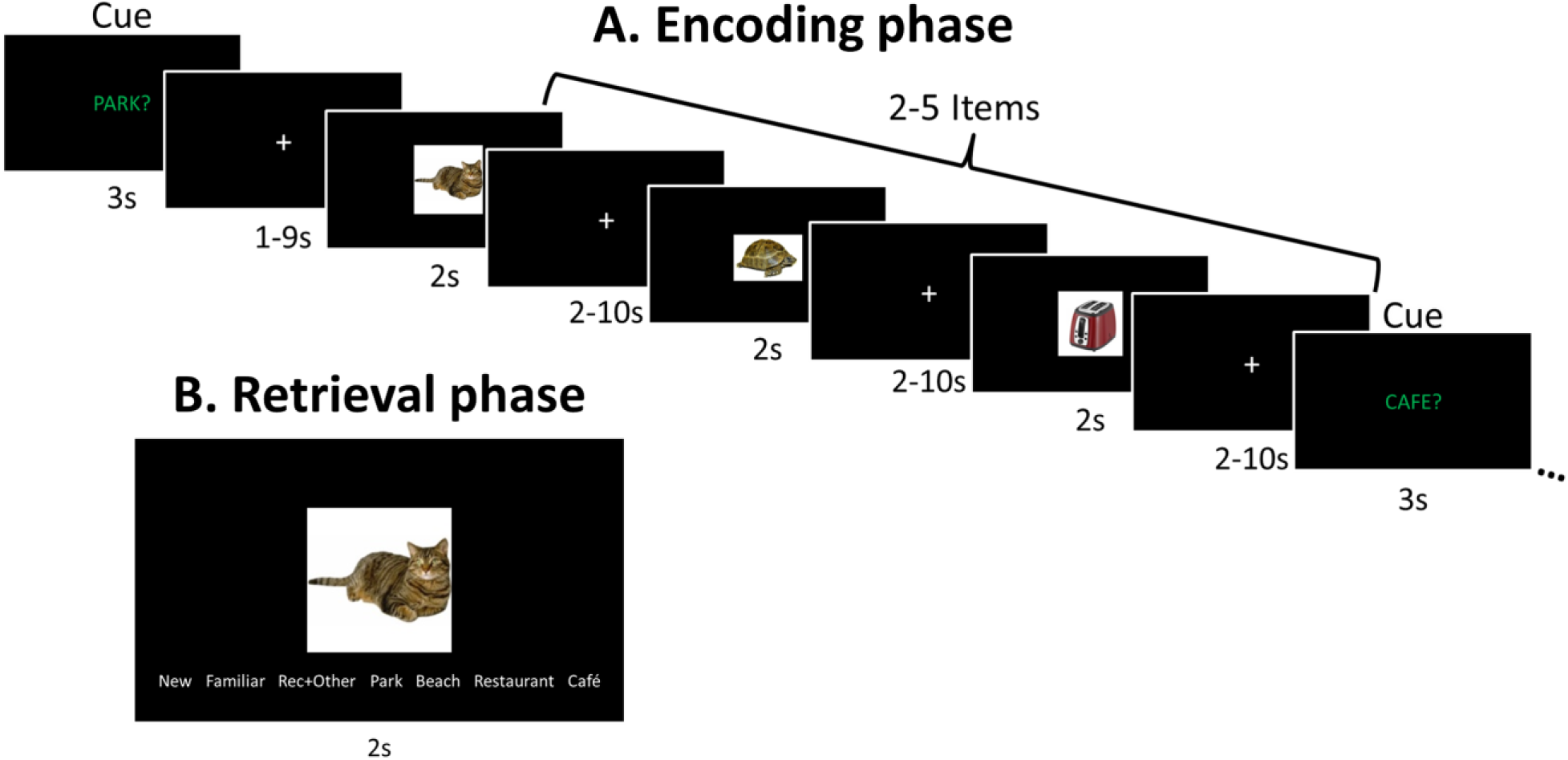
Task design. A) During encoding, trial-unique exemplars from a set of categories were presented sequentially, broken down into mini-blocks separated by a cue screen indicating the encoding context. Two exemplars from each category were encoded in each context. B) At retrieval, all targets and a fully category-matched set of foils were presented in a combined recognition/cued recall test. Only trials given correct source memory judgments were included in voxel pattern similarity analyses.

**Figure 2.**
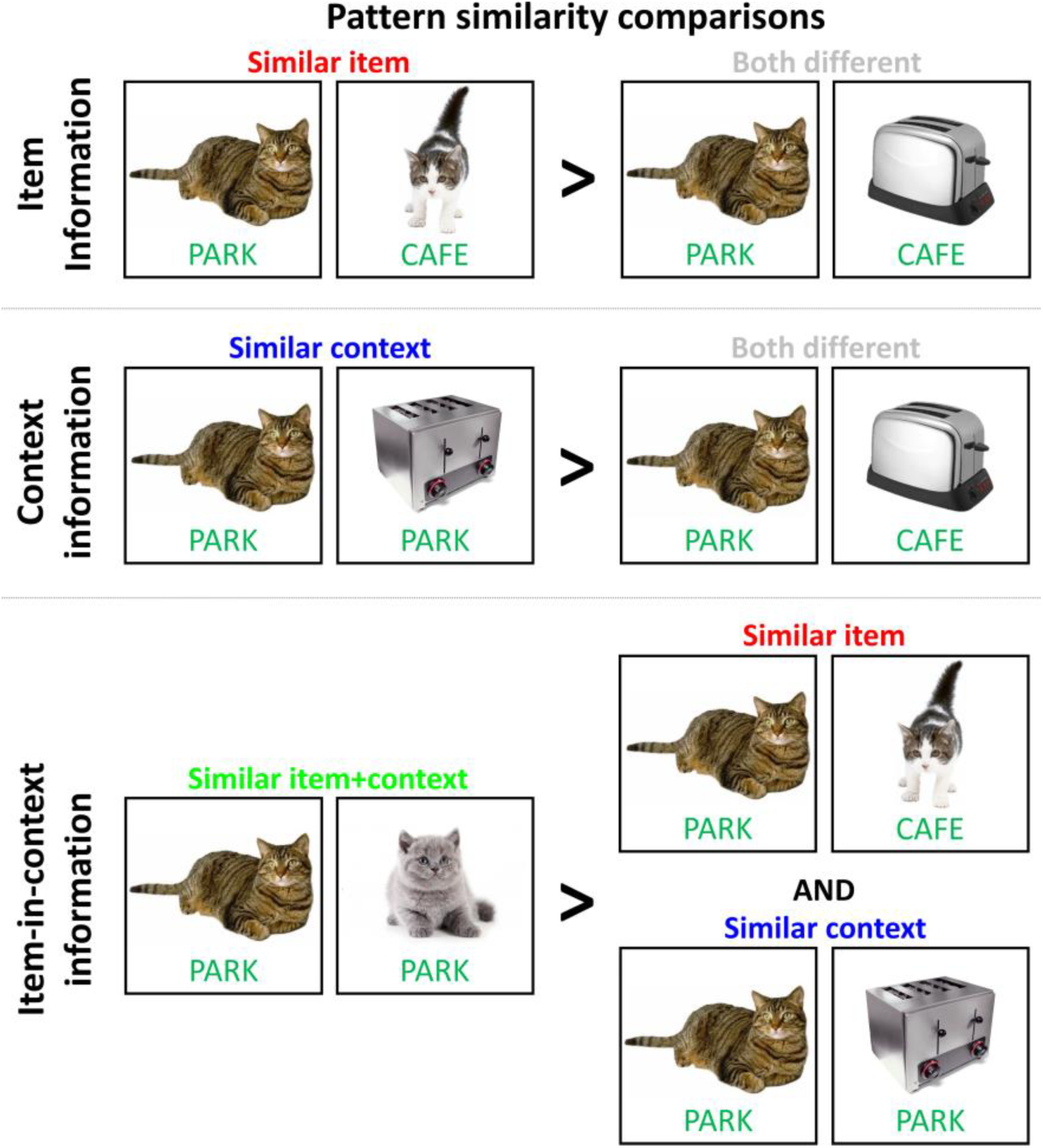
Trial pair similarity predictions. Example trial pair organization for pattern similarity analyses. Item information coding was estimated by contrasting pattern similarity estimates between “similar item” and “both different” trial pairs. Context coding was estimated by contrasting “similar context” and “both different” trial pairs. Item-in-context coding was calculated by contrasting “similar item+context” trial pairs against both “similar item” and “similar context” pairs.

## MATERIALS AND METHODS

### Participants

Twenty-five healthy human participants (female=11; age mean=24.8 years, standard deviation=1.7 years) underwent fMRI scanning during a long-term memory encoding and retrieval task. Participants with head motion greater than 3 mm from origin (N=2) or chance-level context memory performance (N=3) were excluded from analysis, for a total of 20 included participants (female=8). All procedures were approved by the University of California, Davis Institutional Review Board.

### Materials

Item stimuli consisted of 336 visual objects evenly distributed across 28 categories (e.g., toasters, dogs, jackets, muffins) with 12 unique exemplars per category. For each participant, within each category, eight objects were randomly assigned to the list of target stimuli and four objects were assigned to the list of foils, for a total of 224 targets (presented in the encoding and retrieval phases) and 112 foils (presented only in the retrieval phase). Stratified by category, target stimuli were then randomly assigned to one of four encoding context locations (either park, beach, restaurant, or café), for a total of 56 stimuli per context. Therefore, two exemplars from each category were presented in each context.

### Procedure

#### Context personalization and task practice phase – Behavioral

During the encoding phase, participants were cued to visualize each item in one of four spatial contexts: a park, a beach, a restaurant, or a café; at test, participants were asked to retrieve the context originally associated with each item. To ensure that the locations evoked by context cues were visualizable and consistent over the course of the task, context cues were personalized for each participant prior to the start of the task. Participants were asked to select a specific park, beach, restaurant, and café from their past experience that they could visualize well, and were then instructed to bring only those venues to mind in response to the context cues. Participants then received task instructions and completed practice encoding and retrieval runs (using the same four context cues but an independent practice set of item stimuli) on a computer outside of the scanner. Participants were permitted to repeat the practice runs as many times as was necessary to become comfortable with the task timing and response mappings.

#### Encoding phase

Over four encoding runs (Figure 1a), target items were presented sequentially for 2 s each. Item trials were broken down into mini-blocks varying in size from two to five items per block. At the beginning of each mini-block, one of four context cue screens was presented for 3 s, either “PARK,” “BEACH,” “RESTAURANT,” or “CAFÉ,” cuing participants to visualize a specific context location for the duration of the mini-block. Then, for each item in the mini-block, participants visualized the item in that particular context location and made a yes/no response (via a button press) to the question, “In real life, would you be likely to see this item in this location?” Item category, context location, and item category-context pairings were counterbalanced across runs. Within run, item order was pseudo-randomized for each participant, with the constraint that no adjacent item trials contained exemplars from the same category. Block order was also pseudo-randomized for each participant such that no adjacent mini-blocks contained the same context cue, and block length was counterbalanced across context question and across scanning run. Context cue and item trial screens were presented with a jittered ISI optimized for event-related fMRI (SOA 4-12 s with a mean of 6 s).

#### Retrieval phase

Immediately following each encoding run, participants completed a mixed item recognition/source memory test during fMRI scanning (Figure 1b). All target items from the most recent encoding run and a full set of category-matched foils were presented sequentially for 2 s each. For each trial, participants indicated the quality and content of their memory for that item by selecting one of seven response button options: 1) the item was novel (“New”); 2) they were familiar with the item but could not recollect any contextual details associated with having seen it before (“Familiar”); 3) they recollected having seen the item before and could bring some contextual information to mind but *not* the location context in which the item was originally encoded (“Recollect”); 4-7) if they recollected the item and could remember which of the four encoding context locations was originally associated with that item, they pressed a button indicating that location (“Park,” “Beach,” “Restaurant,” or “Café”). Retrieval trials were pseudo-randomized such that no adjacent trials contained exemplars from the same category and jittered for event-related fMRI (SOA=4-12 s with a mean of 6 s). In total, four encoding runs were interleaved with four retrieval runs.

### Image acquisition

MRI scanning was conducted at the UC Davis Facility for Integrative Neuroscience on a 3T Siemens Skyra magnet with a 32-channel phased-array head coil. To facilitate functional image normalization and anatomical localization, high-resolution T1-weighted structural images were acquired using a magnetization-prepared rapid acquisition gradient echo (MPRAGE) pulse sequence (1 mm^3^ voxels; matrix size=256 x 256; 208 slices). Images sensitive to BOLD contrast were acquired using a whole-brain multiband gradient echo planar imaging (EPI) sequence (3 mm^3^ voxels; TR=1220 ms; TE=24 ms; FA=67°; multiband acceleration factor=2; 38 interleaved slices; FOV=192 mm; matrix size=64 x 64) during task performance.

### Behavioral analysis

For the purposes of the fMRI-based multivoxel pattern similarity analysis, detailed below, it was important to establish 1) whether there were any systematic differences in item recognition or context memory across encoding contexts; and 2) that correct context memory judgments were likely to be driven by real memory signal. To measure item discriminability (d’), item hit rate was calculated as the percentage of target trials given any “old” item response (“Familiar,” “Recollect,” or any of the encoding context location judgments – even if the specific location was incorrect) for items encoded in each context location – and overall item false alarm rate was calculated as the proportion of “old” item responses given on foil trials. Differences in item d’ across encoding contexts were tested via a one-way repeated measures ANOVA (with a Huynh-Feldt correction for non-sphericity).

To estimate the meaningfulness of context memory judgments, context discriminability (d’) was calculated separately for each encoding context as the difference between the probability of a correct context judgment (e.g., an item encoded in the café given a “Café” response at test) and the probability of a false alarm to that particular context location (e.g., an item encoded in the park or a foil item given a “Café” response at test). Subjects with context d’ at floor for any encoding context were excluded from fMRI analysis (N=3). Context discriminability estimates were entered into a one-way repeated measures ANOVA to test for differences across encoding contexts (with a Huynh-Feldt correction for non-sphericity).

### Multivoxel pattern similarity analysis

Because we were interested in understanding how the MTL organizes item and context information during successful reinstatement of item-context relations, multivoxel pattern similarity analysis focused on fMRI data from the retrieval phase. EPI timeseries underwent motion correction and highpass filtering (0.01 Hz) in FMRIB’s Software Library (FSL). In preparation for multivoxel pattern similarity analysis procedures, spatial smoothing was omitted. Event-related blood oxygenation level-dependent (BOLD) signal change was estimated separately for each item trial, controlling for signal change due to all other trials and motion artifact, using ordinary least squares regression, resulting 336 single-trial beta images (Mumford, Turner, Ashby, & Poldrack, 2012; Xue et al., 2010). Single-trial beta images from runs 2-4 were coregistered with single-trial beta images from run 1 using FSL’s FLIRT linear registration software (6 degrees of freedom). Coregistered single-trial beta images with atypically high mean absolute z-score (based on the distribution of beta estimates for each grey matter voxel across all trials) were excluded from further analysis. Based on a mean absolute z threshold of 1.5, between 0 and 10 trials were excluded per subject with a median of 4.5 trials. This objective noise trial exclusion procedure has been effective in previous pattern similarity studies (Libby et al., 2014), and, in the current study, beta images were additionally visually inspected to verify that subjectively aberrant trials were not overlooked. Critically, to pinpoint retrieval trials containing both (perceived) item and (reinstated) context information, the subsequent pattern similarity analysis was restricted to correct context memory judgment trials.

To determine the spatial distribution of information coding effects across the hippocampus and surrounding cortical areas, we employed multivoxel pattern similarity analysis (Kriegeskorte, 2011; Kriegeskorte et al., 2008) using a searchlight approach (Kriegeskorte et al., 2006). For every voxel in the brain, correlation coefficients (Pearson’s *r*) were calculated for all pairs of trials based on the pattern of beta coefficients contained in a sphere with a 5-voxel diameter centered on that voxel, resulting in an observed pattern similarity (correlation) matrix. To reduce the influence of temporal autocorrelation and within-run dependencies on pattern information coding effects (Mumford, Davis, & Poldrack, 2014), observed similarity estimates from trial pairs within the same fMRI run were discarded. Between-run observed pattern similarity estimates were entered into second-order similarity analyses. Here, we refer to “second-order similarity” as the z-transformed point-biserial correlation between observed pattern similarity estimates and a binary model of predicted pattern similarity, where trial pairs that were similar along a dimension of interest were coded as 1 and trial pairs that differed along that dimension were coded as 0. In this case, the point-biserial correlation is effectively identical to a two-sample t-test, and was selected because it is sensitive to within-subject variance, providing a somewhat conservative metric of multivoxel pattern similarity.

We examined pattern similarity effects based on shared item and context information in two separate analyses with orthogonal predicted pattern similarity models. To identify brain regions where voxel patterns carried information about the item category, a binary model was constructed labeling trial pairs according to whether they contained items from the same category (e.g., two dog trials – “*similar item*” pairs) or different categories (e.g., one dog trial and one toaster trial – “*both different*” pairs), excluding trial pairs with the same context location. To identify brain regions with voxel patterns carrying retrieved context information, a binary model was constructed labeling trial pairs according to whether the retrieved context was the same (e.g., two park trials – “*similar context*” pairs) or different (e.g., a park trial and a beach trial – “*both different*” pairs), excluding trial pairs with the same item category.

To test for pattern similarity effects driven by shared item-context relations that were distinguishable from item or context effects alone, we constructed a binary model for item-in-context similarity that was superordinate to the item and context models (Figure 2, Supplementary Figure 1). Trial pairs with both item category and context location in common (e.g., two dog-in-the-park trials – “*similar item+context*” pairs) were coded as 1 and trial pairs with overlapping item category *or* context location, but not both (i.e. *similar item* and *similar context* pairs), were coded as 0; trial pairs that were different on both item category and context location dimensions were excluded.

For each of the three predicted similarity models, the resulting second-order similarity estimate was assigned to the center voxel of each searchlight, resulting in three whole-brain pattern similarity images for each subject. Single-subject pattern similarity images were normalized to the MNI152 template for group analysis via a two-step registration process. Using FLIRT, representative single-subject EPI images were co-registered with high-resolution anatomical MPRAGE images (6 degrees of freedom), MPRAGE images were normalized to the template image (12 degrees of freedom), and the two resulting transformation matrices were concatenated; the resulting EPI-to-template linear transformation matrices were then applied to the pattern similarity images.

Brain regions that reliably carried each type of information across subjects (*p*_FWE_ < 0.05) were identified non-parametrically based on the distribution of maximum cluster mass (after a voxelwise t-threshold of 2.539, or *p* < 0.01) across 10,000 permutations of the data (sign-flipping approach) using the Randomise function in FSL (Nichols & Holmes, 2002). Because we had *a priori* interest in the hippocampus and surrounding cortex, group analyses were restricted to MTL voxels using a statistically conservative (anatomically liberal) MTL mask generated with the WFU Pickatlas (Maldjian, Laurienti, Kraft, & Burdette, 2003) consisting of parahippocampal gyrus, hippocampus, uncus, and amygdala and dilated in three dimensions by 4 mm. The dilated anatomical MTL mask extended laterally and ventrally into fusiform cortex, posteriorly to the level of the posterior horn of the lateral ventricles, and anteriorly into medial temporopolar cortex. We additionally explored pattern similarity effects outside of the MTL using the same non-parametric permutation test procedure, but with whole-brain grey matter masks (p(grey matter) > 0.36, corresponding to a threshold of 90 out of 245 of the FSL MNI 152 segmentation grey matter priors) that excluded voxels contained in the MTL mask.

Because the above one-sample t-tests did not yield significant voxels where pattern similarity related to context similarity (independent of item similarity) on average across subjects, we hypothesized that the strength of neural pattern similarity evidence for context reinstatement might be related to individual differences in behavioral evidence for context memory. To test this idea, in a voxelwise regression (with non-parametric inference testing, as above, but with full permutations instead of sign-flipping), we identified regions where pattern similarity evidence related to context similarity were positively associated with individual behavioral context discriminability estimates (averaged across contexts). Family-wise error rate was determined on the basis of cluster mass using MTL and non-MTL grey matter masks, as described above. However, no regions in the MTL surpassed the stringent cluster mass-based family-wise error rate correction applied in other analyses. Exploratory thresholding was then applied to the MTL results, using a relaxed voxelwise threshold of *p* < 0.05 and a cluster mass threshold set to p < 0.05, FWE-corrected.

To allow for visualization and comparison of pattern similarity effects specifically in the hippocampus, for each subject, a second set of searchlight images was generated containing average pattern similarity estimates for each combination of item category and context location overlap. That is, one image contained average similarity estimates for *similar item* pairs, one image contained average similarity estimates for *similar context* pairs, one image contained the average across *similar item+context* pairs, and one image contained the average across trial pairs that were different on both item category and context location (“*both different*” pairs). From these images, mean pattern similarity estimates were extracted using individually-defined regions of interest (ROIs) corresponding to head, body, and tail divisions of the hippocampus. Because items-in-context and pattern separation accounts of MTL function both predict that adjacent MTL cortical areas tend to generalize across similar events, pattern similarity estimates were also extracted from anatomical ROIs in perirhinal cortex and parahippocampal cortex as a basis of comparison to any hippocampal generalization effects. All ROIs were defined separately for right and left hemispheres, and anatomical landmarks for ROI definition are described nicely in (Moore et al., 2014).

Pattern similarity estimates were fed into a full factorial repeated measures ANOVA with ROI and information overlap as factors of interest (using a Huynh-Feldt correction for non-sphericity), controlling for hemisphere. Significant interaction effects were broken down separately by ROI. *A priori* contrasts testing for item generalization (*similar item* > *both different*), context generalization (*similar context* > *both different*), and item-in-context generalization (*similar item+context* > *similar item+similar context*) were defined for information overlap, in keeping with the prediction model hierarchy described above. When two or more *a priori* contrasts were significant within an ROI, we also applied *post hoc* contrasts (with Bonferroni correction for multiple comparisons) to determine which significant effect was strongest (e.g., *similar item+context* > *similar item*). Contrasts were additionally applied to pattern similarity estimates extracted from eroded hippocampal head ROIs that excluded edge voxels (binary erosion in three dimensions with 3mm spherical kernel) to reduce any possible influence of signal from neighboring voxels.

Finally, we conducted a control analysis to determine whether pattern similarity in hippocampal head could be driven by the similarity of motor responses between trials, independent of memory for context information. Evoked patterns of activity across voxels in hippocampal head ROIs were correlated for each pair of trials given either a “Familiar” response (trials characterized by item familiarity without context recollection) or “New” response (forgotten trials). If pattern similarity was merely sensitive to motor response rather than episodic memory details, similarity estimates between trial pairs containing two “Familiar” or two “New” trials (i.e., *within* trials with the same motor response) would be expected to be higher than between trial pairs containing one “Familiar” and one “New” trial (i.e., *between* trials with different motor responses). Mean similarity estimates were entered into a response similarity (*within familiar*, *within forgotten*, *between*) repeated measures ANOVA, controlling for hemisphere.

## RESULTS

Because we were interested in voxel patterns associated with memory for items in context, it was important to establish the presence of two behavioral effects: 1) that memory (particularly context memory) *existed*, i.e. that any given correct context memory response could be attributed to real memory signal; and 2) that there were no systematic differences in memory performance across encoding conditions. One-way repeated measures ANOVAs confirmed that both item discriminability (F_(1,19)_ = 238.34, *p* < 0.0001) and context discriminability (F_(1,19)_ = 77.20, *p* < 0.0001) were significantly greater than zero across encoding contexts, but that neither metric differed significantly between encoding contexts (F’s < 1.39, *p*’s > 0.25). A summary of these behavioral effects for each context location is provided in Supplementary Figure 2.

To examine how the hippocampus organizes memories for similar events, we performed a set of multivoxel pattern similarity searchlight analyses (Kriegeskorte et al., 2006; Kriegeskorte et al., 2008). Analysis was restricted to retrieval phase trials with accurate context recall decisions. Local voxelwise patterns of evoked activity were correlated between all pairs of these trials, and trial pair correlations were contrasted according to the similarity of item and/or context information contained in each pair (Figure 2, Supplementary Figure 1).

We first investigated whether local voxel patterns in the hippocampus generalized across different exemplars of the same item. Voxelwise pattern similarity was contrasted between trial pairs containing different item exemplars from the same category (“*similar item”* pairs) against pattern similarity between trial pairs that included exemplars from different item categories (“*both different”* pairs). To isolate item-level pattern information, this analysis excluded trial pairs that had been associated with the same encoding context. Within a medial temporal lobe (MTL) mask, significant clusters extended throughout the ventral and medial temporal neocortex, including perirhinal cortex, and amygdala, but no hippocampal voxels were identified as generalizing across events on the basis of item information alone (Supplementary Figure 3a), consistent with previous studies of item pattern separation in the hippocampus and medial temporal cortex (Huffman & Stark, 2014; LaRocque et al., 2013; Liang et al., 2012). Exploratory whole brain analyses excluding MTL voxels revealed item-level pattern information in frontal, parietal, and occipital regions, with the strongest effects in right lateral occipital cortex and posterior fusiform cortex (Supplementary Figure 3b).

We next examined whether the hippocampus generalized across different items according to whether they had been studied with the same encoding context. We contrasted voxel pattern similarity between trial pairs with different item cues that shared a study context (“*similar context”* pairs) and *both different* pairs. No suprathreshold clusters in hippocampus or neocortex were identified as showing higher pattern similarity for similar context than for both different pairs, either within the MTL or across the whole brain. We next investigated whether neural evidence for context reinstatement was related to individual differences in the behavioral expression of context memory. This analysis revealed regions in parahippocampal cortex (MTL mask), as well as retrosplenial cortex, lingual gyrus, and precuneus (exploratory whole-brain mask), where pattern similarity related to context information (independent of item information) was positively correlated with context memory discriminability across subjects (Supplementary Figure 4). However, again, no significant hippocampal voxels were identified.

Our third analysis tested the possibility that the hippocampus generalized across memories that included similar item <u>and</u> context information. We predicted that, if the hippocampus organized memories on the basis of item-context associations, voxel pattern similarity would be substantially higher across events with similar items and similar contexts, compared to events that shared only a single attribute. Therefore, we contrasted voxel pattern similarity between trial pairs with that included similar items that had been associated with the same study context location (“*similar item+context”* pairs) against pattern similarity between trial pairs that overlapped on one, but not both, of these dimensions (i.e., *similar item* and *similar context* pairs). Searchlight analysis identified clusters showing this effect with centers of mass in the anterior and middle hippocampus bilaterally, extending partially into ventral and medial temporal neocortex (Figure 3a,b).

**Figure 3.**
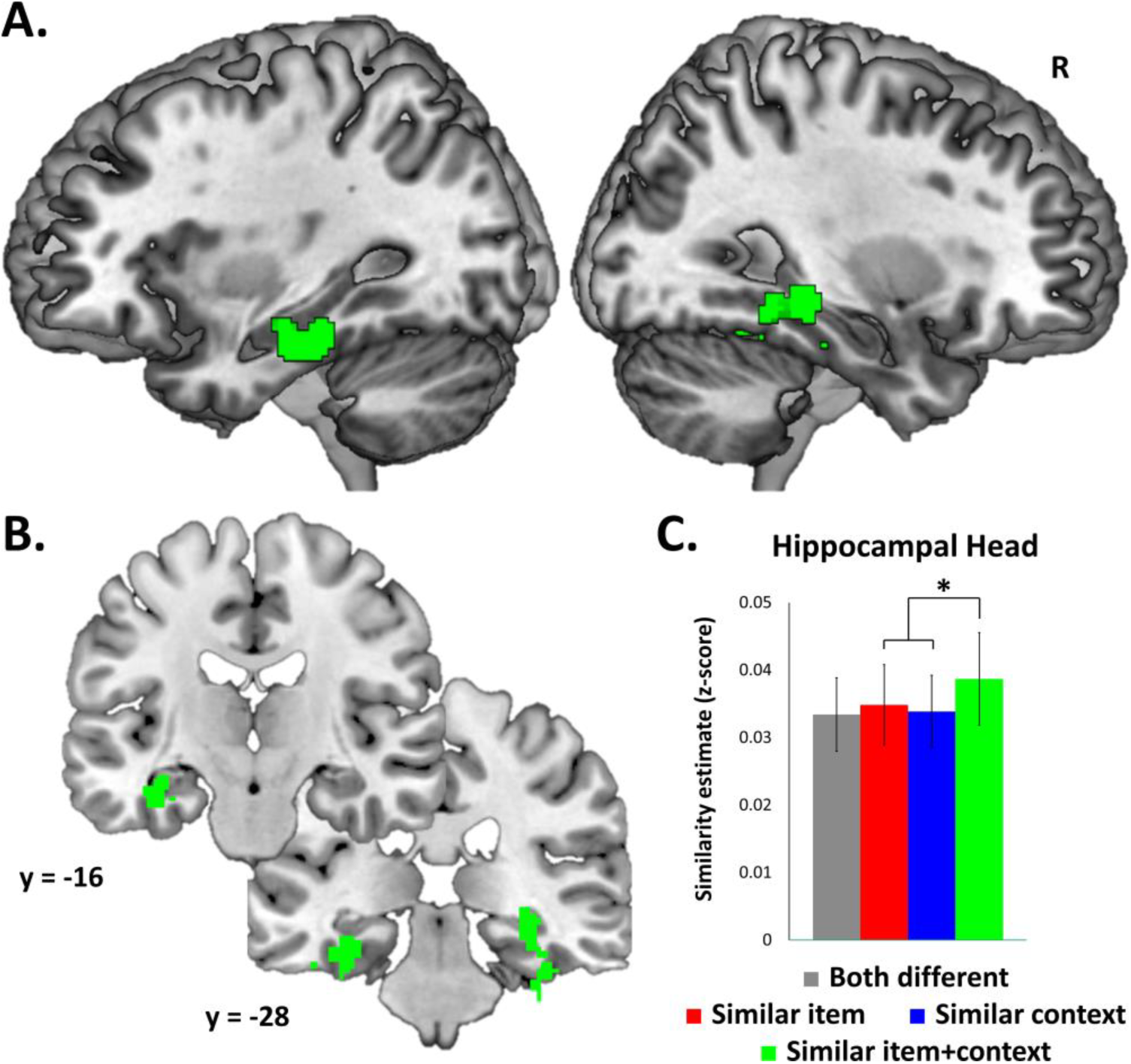
Hippocampal voxel pattern similarity analysis results. A-B) Searchlight results for similar item+context > (similar item or similar context) contrast in sagittal (A) and coronal (B) views (voxelwise threshold p < 0.01, FWE-corrected cluster p < 0.05). C) Mean pattern similarity estimates for both different, similar item, similar context, and similar item+context trial pairs extracted from anatomical hippocampal head ROIs. Because there was no significant difference in the pattern of effects by hemisphere, results are collapsed across right and left ROIs. Error bars represent standard error of the mean. * *p* < 0.05.

To further characterize item and context organization within the hippocampus and adjacent cortical regions, mean pattern similarity estimates (z-scores) for *both different*, *similar item*, *similar context*, and *similar item+context* pairs were extracted from anatomically-defined hippocampal and cortical regions of interest (ROIs). Because of previously reported functional differences along the anterior-posterior axes of the hippocampus (Poppenk, Evensmoen, Moscovitch, & Nadel, 2013), we evaluated hippocampal head, body, and tail separately using established anatomical landmarks (Moore et al., 2014). A full factorial repeated measures ANOVA revealed a significant main effect of ROI (F_(4,19)_=16.37, corrected *p*_HF_ < 0.0001, ε_HF_=0.54), a significant main effect of information overlap (F_(3,19)_=4.45, corrected *p*_HF_ < 0.05, ε_HF_=0.41), and, critically, a significant ROI-by-information overlap interaction (F_(12,19)_=4.25, corrected *p*_HF_ < 0.005, εHF=0.36); there was no significant main effect of hemisphere or interaction between hemisphere and either factor of interest (F’s < 1.58, *p*’s > 0.18). Based on the interaction effect, we examined the effect of information overlap within each ROI, controlling for hemisphere, using a set of *a priori* and follow-up *post hoc* contrasts to detect the presence of item, context, and item-in-context generalization separately for each ROI.

Information generalization effects in medial temporal lobe cortical ROIs were examined as a basis of comparison for hippocampus (Supplementary Figure 5). Within perirhinal cortex, pattern similarity estimates were greater for *similar item* compared to *both different* trial pairs (F_(1,19)_=3.12, *p*=0.08, trend-level), and also greater for *similar item+context* relative to *similar item* pairs (F_(1,19)_=12.9, *p* < 0.005, Bonferroni-corrected), suggesting that item generalization in perirhinal cortex is strengthened by context similarity. In parahippocampal cortex, pattern similarity between *similar item* pairs was significantly greater than *both different* pairs (F_(1,19)_=8.38, *p* < 0.01), but were not significantly different from *similar item+context* pairs (F_(1,19)_=12.9, *p* < 0.005, Bonferroni-corrected), suggesting that item generalization in parahippocampal cortex was insensitive to context similarity.

In contrast to item generalization effects in the cortex, within hippocampal head, pattern similarity estimates were significantly and selectively greater for *similar item+context* pairs compared to *similar item* and *similar context* pairs (F_(1,19)_=5.19, *p* < 0.05), consistent with voxelwise results (Figure 3c). This pattern of results suggests that hippocampal head only generalized across trials that shared similar item-in-context information. Eroded hippocampal head ROIs excluding edge voxels demonstrated a qualitatively similar pattern of results, although the selective similar item+context effect was attenuated (F_(1,19)_=3.68, *p*=0.06), suggesting this effect was not driven by adjacent non-hippocampal signal. There were no significant pattern similarity differences in hippocampal body or tail ROIs (F’s < 1.72, *p*’s > 0.19).

Finally, to test for the influence of motor response similarity on the selective *similar item+context* effect in the hippocampal head, pattern similarity was contrasted between incorrect trial pairs that were associated with the same button press against incorrect trials that were associated with different button presses (“incorrect” here referring to trials not reflecting item-in-context memory). If hippocampal patterns were driven by motor responses alone, rather than the content of episodic memories, pattern similarity would be expected to be higher between two trials endorsed as familiar, as well as between two trials that were forgotten, compared to similarity between one familiar trial and one forgotten trial. However, a repeated measures ANOVA on response similarity, controlling for hemisphere, revealed no significant effects (F’s < 1.89, p’s > 0.18), suggesting that hippocampal pattern similarity was not driven by specific motor responses.

## DISCUSSION

In the current study, we used multivoxel pattern similarity analysis to investigate event representation in the hippocampus and MTL cortical areas during episodic retrieval. Anterior hippocampal voxel activity patterns were similar for events that involved similar items that had been associated with the same study context, but not for events with similar item or context information in isolation. These findings provide novel insight into the neural basis of memory for episodic details, suggesting that, although neural coding in the hippocampus may differentiate between events with some overlapping attributes, it generalizes across events that share similar item-context relations.

Several recent human fMRI studies (Bakker, Kirwan, Miller, & Stark, 2008; Huffman & Stark, 2014; Lacy, Yassa, Stark, Muftuler, & Stark, 2011; LaRocque et al., 2013; Liang et al., 2012) have shown that, compared to MTL cortical regions, the hippocampus (particularly the dentate gyrus) differentiates between similar items. In the current study, we also found that the hippocampus did not generalize across similar items, as long as they had different contextual associations. However, when similar items shared context information, hippocampal activity patterns overlapped. This finding suggests an important boundary condition on hippocampal pattern separation: the extent to which the hippocampus differentiates between similar items will depend on the context in which the items had been encountered. It is conceivable that imaging with higher resolution could reveal more pronounced pattern separation in specific hippocampal subfields, such as the dentate gyrus (Leutgeb et al., 2007; Rolls & Kesner, 2006; Yassa & Stark, 2011), but previous multivariate pattern similarity studies using high-resolution fMRI have not reported reliable subfield-level differences in item pattern separation (Huffman & Stark, 2014; LaRocque et al., 2013; Liang et al., 2012).

The current results add importantly to previous work on hippocampal coding of item and context information. Although previous studies of hippocampal activity pattern information have not manipulated the similarity of item and context information, these studies have consistently suggested that hippocampal representations of items are context-dependent. Consistent with our results, neuronal ensemble recordings in rats (Manns & Eichenbaum, 2009; McKenzie et al., 2014) have shown that the stability of hippocampal coding for specific items across repeated exposures is dependent on spatial context. Also in line with present findings, a previous fMRI study has shown that voxel patterns in the anterior hippocampus were more consistent across pairs of trials characterized by successful reinstatement of bound item-in-context information, compared to trials without associative retrieval (Hannula, Libby, Yonelinas, & Ranganath, 2013). The current study extends our knowledge by focusing only on trials characterized by associative retrieval, demonstrating that item-in-context coding in the hippocampus is not merely a function of memory quality and is sensitive to the content of individual episodic memories. Additionally, our findings accord with recent fMRI studies in domains outside of episodic memory that relate the fidelity of hippocampal voxel patterns to implicit memory for the temporal position of items in learned sequences (Hsieh et al., 2014) and to accurate working memory for spatial configurations of items (Libby et al., 2014). Together, these results suggest that the hippocampus generally supports cognition by tracking conjunctions of items and their temporal or spatial contexts.

It is noteworthy that we observed similarity in hippocampal voxel patterns across *similar item+context* trial pairs, despite the fact that the corresponding study events were not exact duplicates. The events were similar at a schematic level but differed in their exact features. It could be argued that our results accord with the Complementary Learning Systems (CLS; Norman, 2010; Norman & O’Reilly, 2003) computational model, which predicts that hippocampal pattern separation breaks down when the average overlap across events is high. According to this view, however, we would expect participants to exhibit poor discriminability between related events (Elfman, Parks, & Yonelinas, 2008; Norman, 2010; Norman & O’Reilly, 2003), whereas item and context discriminability were very high in the current study. High memory performance cannot be explained in terms of schema-based retrieval (e.g., “I remember that there was a dog in the park”), because similar exemplars from each category were encoded in each context, and these items had to be discriminated from similar unstudied exemplars. As a result, participants had to use detailed representations to discriminate between related events. The present results therefore suggest that models like CLS can be improved by explicitly incorporating representations of item and context information. If the model were instantiated such that context features play a predominant role in hippocampal representations, then we would expect that the CLS model would tend to exhibit pattern completion during processing of similar items encountered in the similar context, and pattern separation during processing of similar items encountered in different contexts.

Several recent studies have reported a role for the hippocampus in the integration of separate events that share key conceptual information (Chadwick, Hassabis, & Maguire, 2011; Horner, Bisby, Bush, Lin, & Burgess, 2015; Milivojevic, Vicente-Grabovetsky, & Doeller, 2015; Schlichting, Mumford, & Preston, 2015). For instance, Milivojevic and colleagues (2015) showed that hippocampal voxel patterns are more similar across seemingly unrelated narrative videos when an intervening video reveals a link between the people (items) and locations (contexts) in the videos. On the basis of the current results, we would predict that an intervening video that shares context or items with the surrounding episodes, but not both, would disrupt hippocampal integration, perhaps particularly in anterior hippocampus. Indeed, in studies of complex pairwise item associations learned over the course of multiple trials, activity (Horner et al., 2015) and pattern similarity (Schlichting et al., 2015) in anterior hippocampus were greatest when event elements were fully integrated into “holistic” representations after learning. Given that the present study investigated single-shot episodic encoding and retrieval, our results suggest that anterior hippocampus integration or generalization can occur rapidly when item-context relations are sufficiently conceptually overlapping.

Everyday experiences often involve many common elements, yet we are able to retrieve specific instances of item-context associations – to which rack you locked your bike this afternoon, which salad you ate at your desk on Monday, which dog you saw in the park over the weekend – with surprising accuracy (Johnson et al., 1993; Mitchell & Johnson, 2009). Results from the current study suggest that the hippocampus utilizes context as an organizing principle in order to reduce competition between memories that involve the same or similar items in different contexts. These findings bridge two influential theoretical frameworks on hippocampal function (Davachi, 2006; Eacott & Gaffan, 2005; Knierim et al., 2006; Marr, 1971; Norman, 2010; Norman & O’Reilly, 2003; Ranganath, 2010; Ranganath & Ritchey, 2012; Ritchey, Libby, et al., 2015; Rolls & Kesner, 2006; Yassa & Stark, 2011), by suggesting a critical role for the binding of item and context information in hippocampal pattern separation and episodic memory.

## ACKNOWLEDGEMENTS

This work was supported by a National Security Science and Engineering Faculty Fellowship (NSSEFF) from the U.S. Department of Defense, a Guggenheim Fellowship, and the NSF Graduate Research Fellowship Program. In memory of Dr. Michael H. Buonocore.

---

SUPPLEMENTARY FIGURES AND CAPTIONS

**Supplementary Figure 1.**
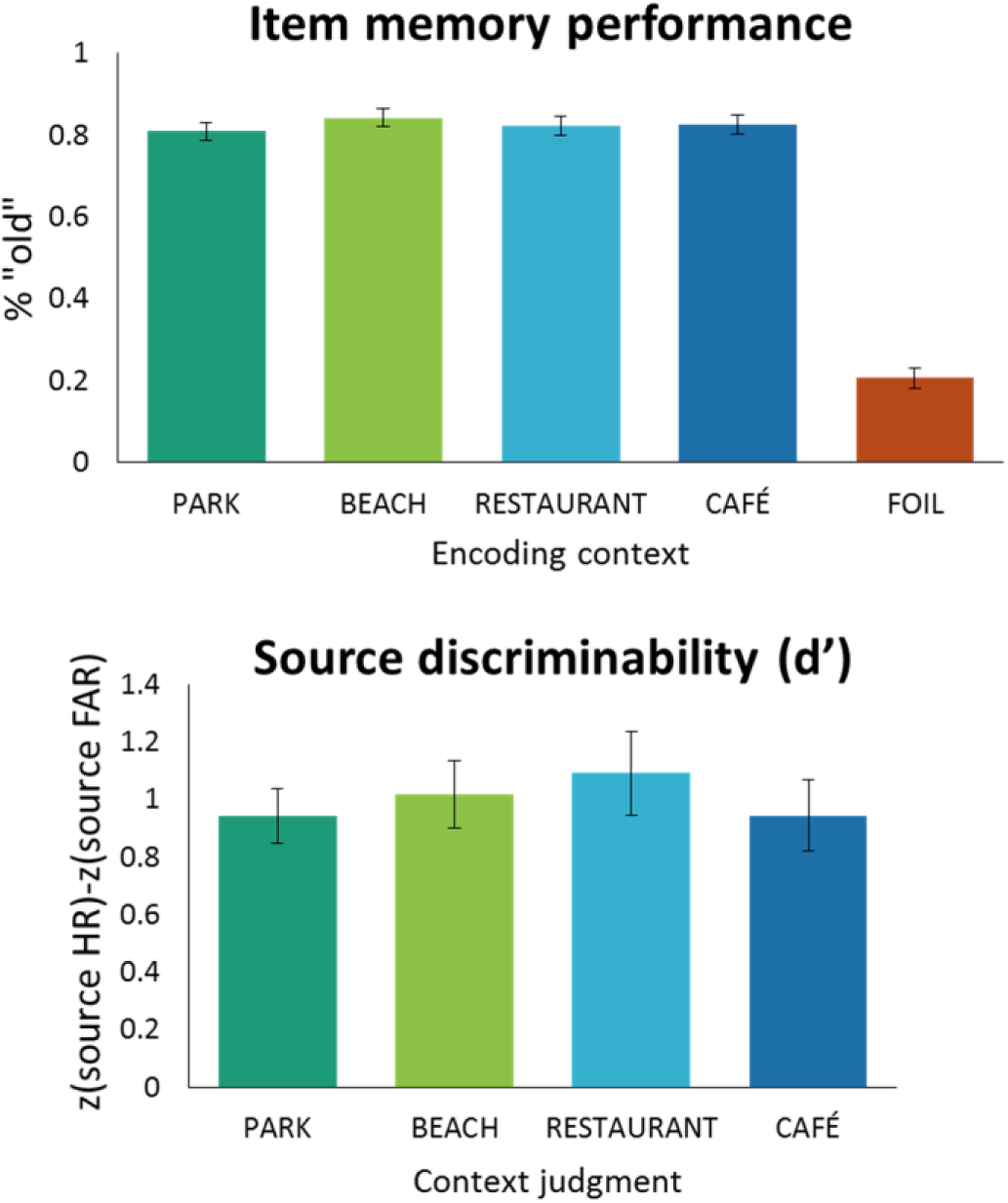
Behavioral expressions of item and context memory were equivalent across encoding context locations. Item memory performance (top) was calculated as the proportion of old items from each encoding context given any “old” response (“Familiar,” “Recollect,” or any source judgment); item false alarm rate was calculated as the proportion of foils given any “old” response. Source discriminability (bottom) was calculated as the difference between the inverse normal cumulative distributions of source hit rate (HR) and source false alarm rate (FAR). For each context judgment, source HR was calculated as the proportion of items correctly attributed to a particular encoding condition; source FAR was calculated as the proportion of items (targets or foils) incorrectly attributed to that encoding condition. Error bars indicate standard error of the mean across participants.

**Supplementary Figure 2.**
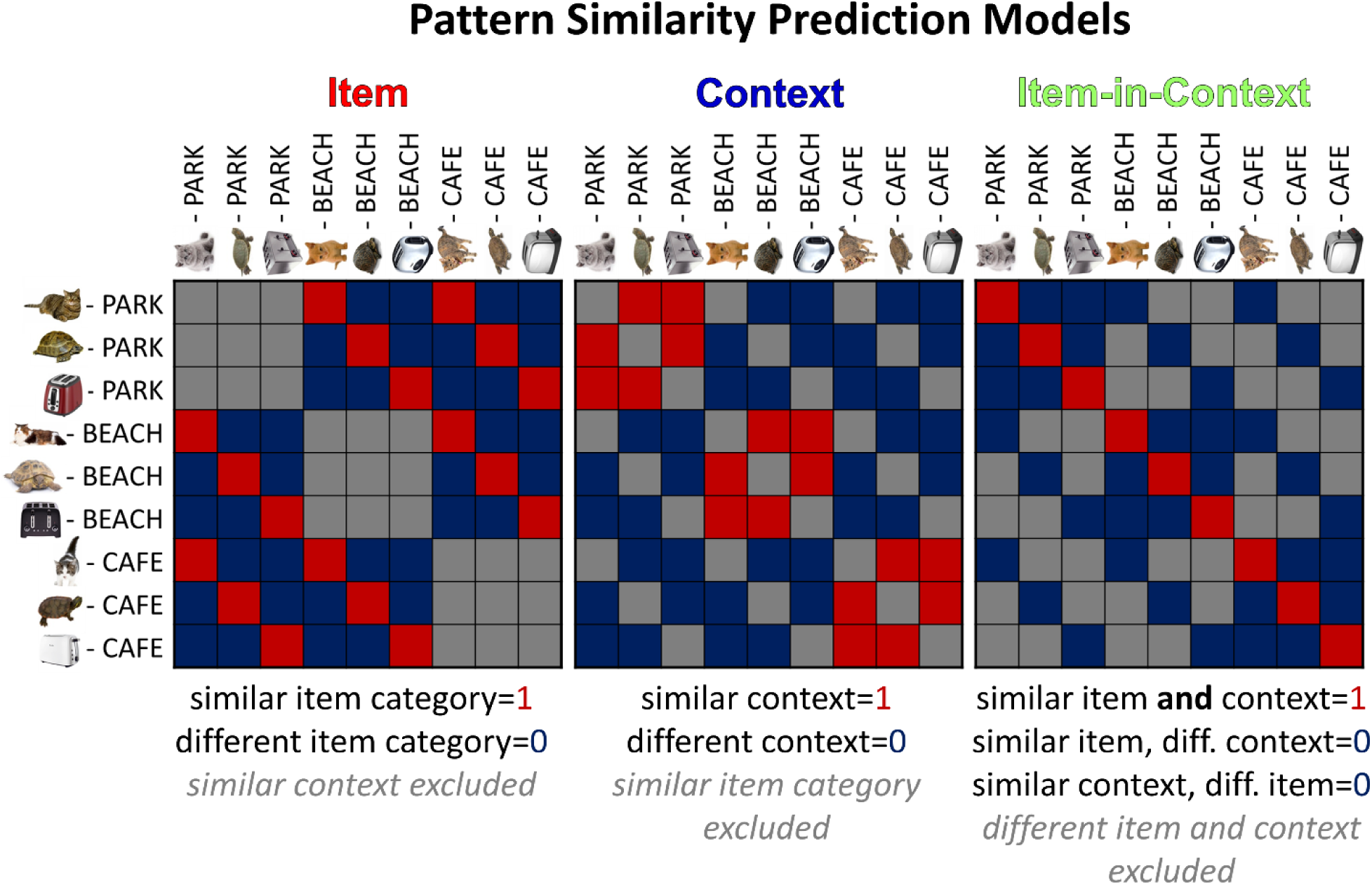
Example segment of predicted trial-by-trial similarity model matrix. Item and context predicted similarity models were orthogonal to each other; the item-in-context predicted similarity model was hierarchical to the item and context models.

**Supplementary Figure 3.**
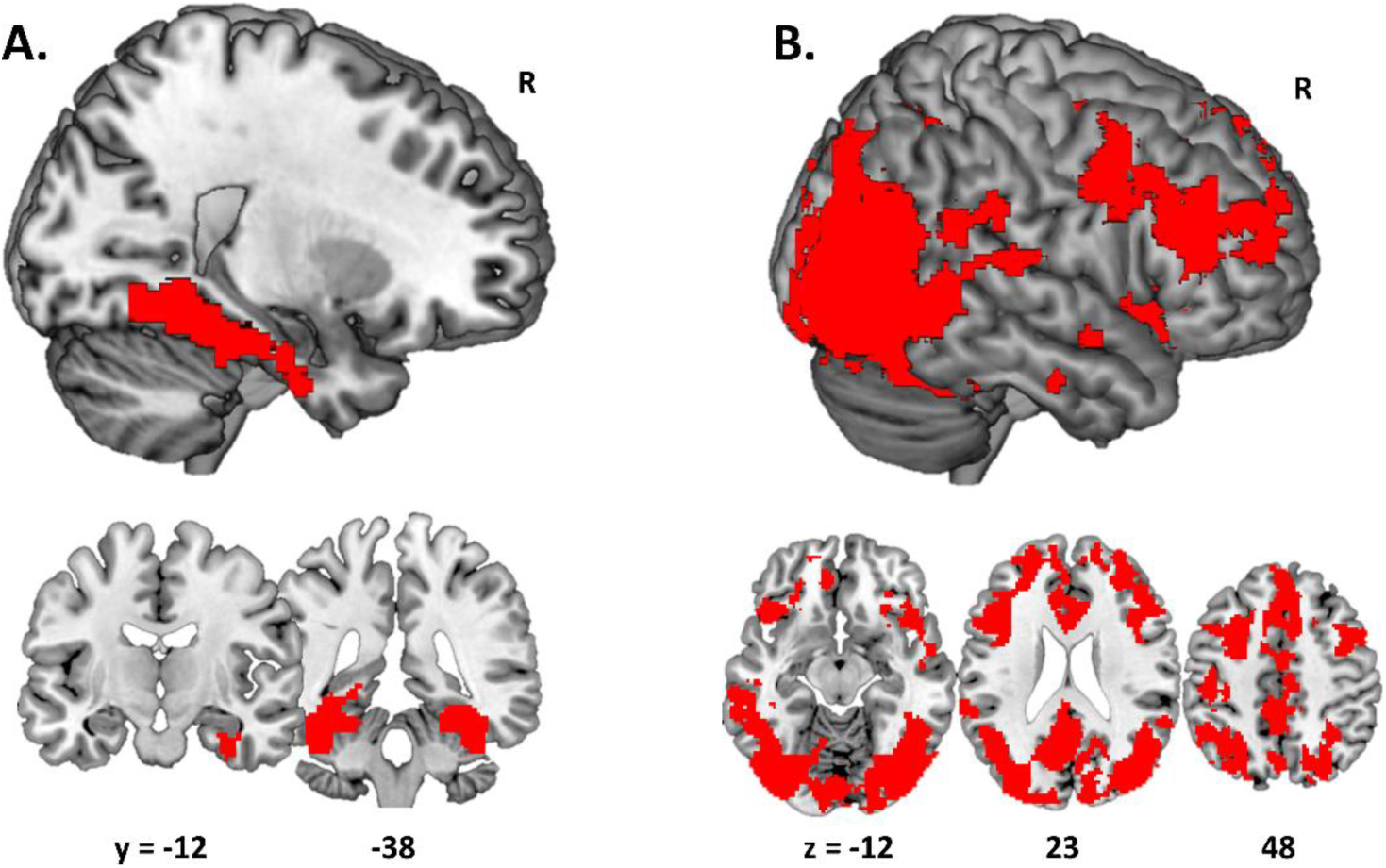
Voxel pattern similarity analysis results for *similar item* > *both different* contrast. A) Using an MTL mask, significant clusters were identified in ventral and medial temporal cortex, including perirhinal cortex. B) Using a whole-brain grey matter mask (excluding MTL voxels), extensive significant clusters were identified, with the global maximum in posterior fusiform cortex (MNI 40, −56, −14; t = 10.22). Voxelwise threshold set to p < 0.01 with FWE-corrected cluster p < 0.05.

**Supplementary Figure 4.**
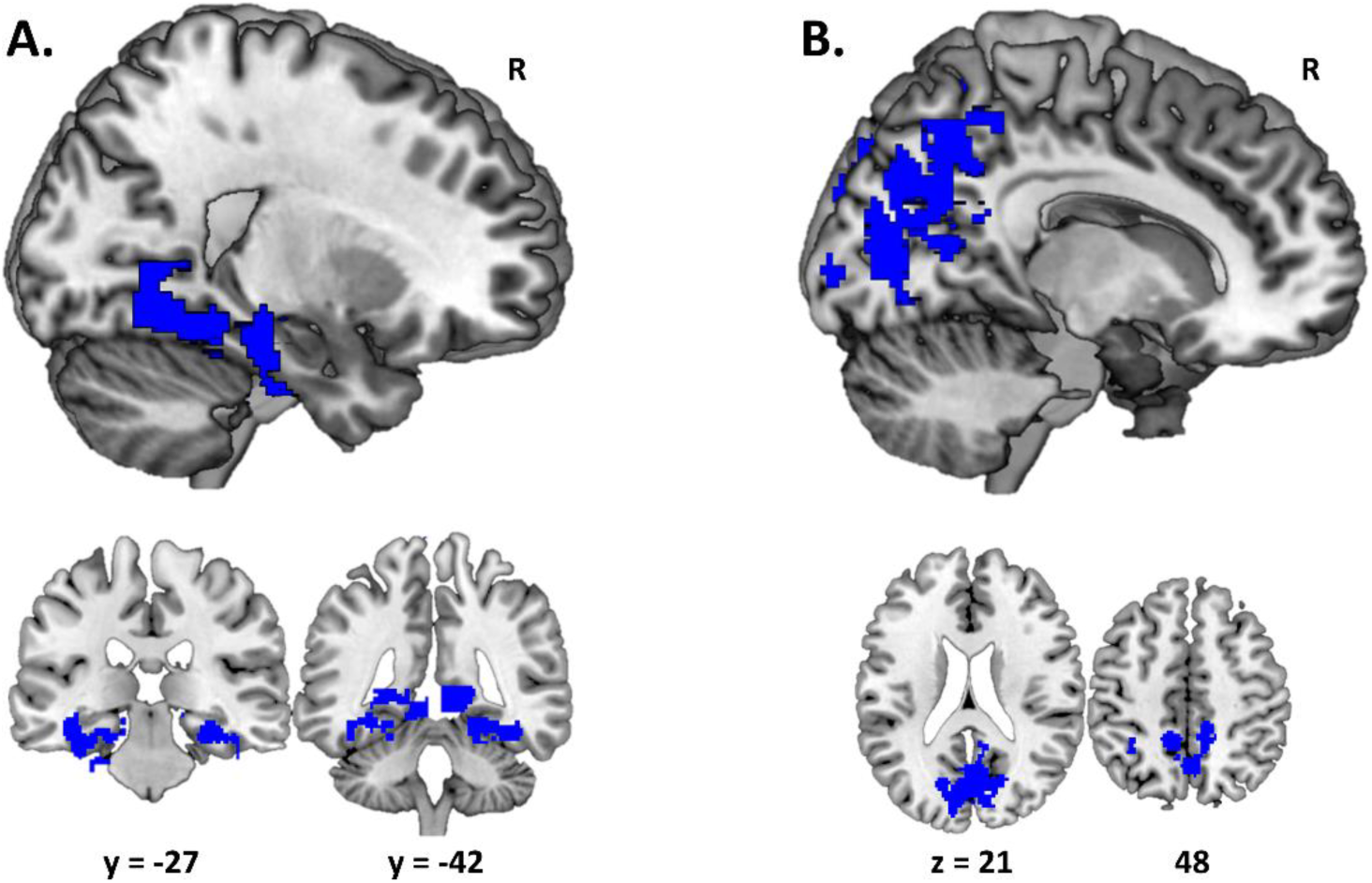
Association between similar context > both different pattern similarity and context memory performance. A) With an MTL mask, significant clusters were identified in posterior-medial temporal cortex, with the global maximum located in parahippocampal cortex (MNI 28, −24, −20; t=6.61). For display purposes, exploratory voxelwise threshold set to p < 0.05 with FWE-corrected cluster p < 0.05. B) With a whole-brain grey matter mask (excluding MTL voxels), significant clusters were identified in posterior medial regions, with the global maximum in cuneus (MNI −10, −86, 16; t = 6.82). Voxelwise threshold set to p < 0.01 with FWE-corrected cluster p < 0.05.

**Supplementary Figure 5.**
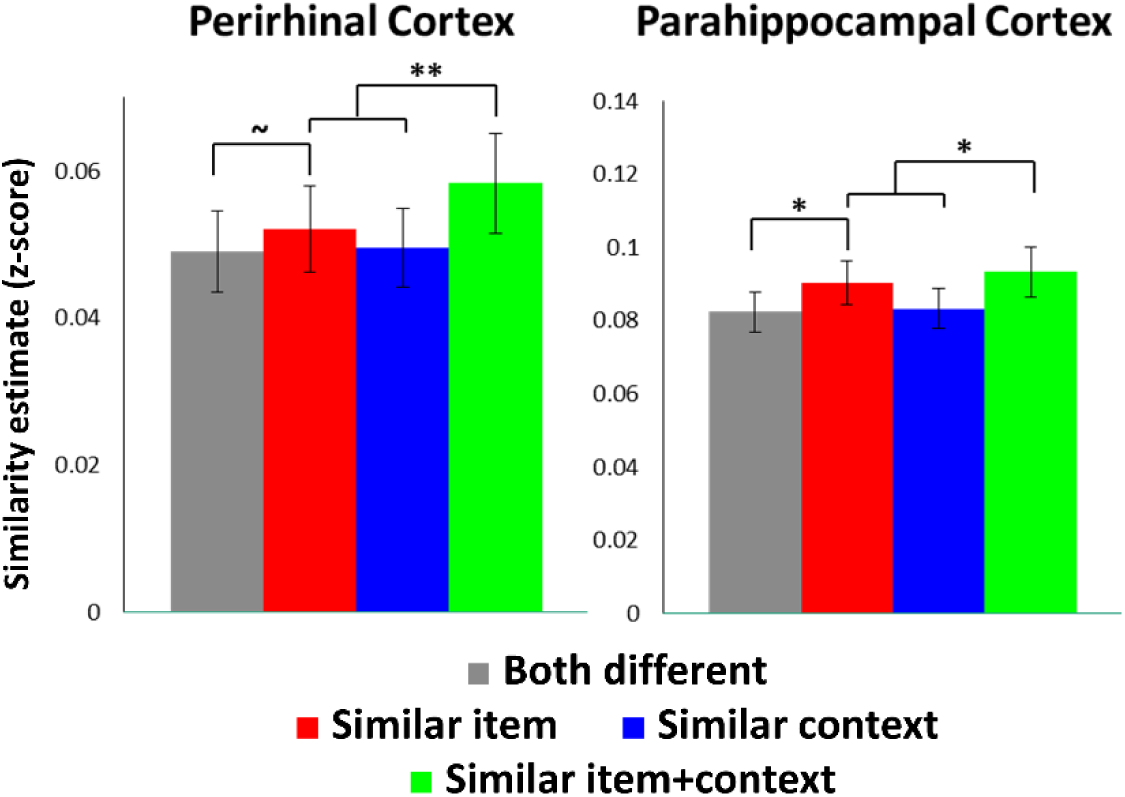
Mean pattern similarity estimates extracted from anatomical MTL cortex ROIs. In PRC, there was evidence for both item and item-in-context generalization. In PHC, item-in-context generalization was redundant with item coding effects. ~ p ≈ 0.05 (trend-level), * p < 0.05 for a priori contrasts (indicated by brackets). ** p < 0.05 for both a priori and post-hoc (similar item+context > similar item) contrasts. Error bars refer to standard error across participants.

## REFERENCES

Bakker, A., Kirwan, C. B., Miller, M., & Stark, C. E. (2008). Pattern separation in the human hippocampal CA3 and dentate gyrus. Science, 319(5870), 1640–1642. doi:319/5870/1640 [pii] 10.1126/science.1152882

Bostock, E., Muller, R. U., & Kubie, J. L. (1991). Experience-dependent modifications of hippocampal place cell firing. Hippocampus, 1(2), 193–205. doi:10.1002/hipo.450010207

Brown, M. W., & Aggleton, J. P. (2001). Recognition memory: what are the roles of the perirhinal cortex and hippocampus? Nat Rev Neurosci, 2(1), 51–61. doi:10.1038/35049064

Chadwick, M. J., Hassabis, D., & Maguire, E. A. (2011). Decoding overlapping memories in the medial temporal lobes using high-resolution fMRI. Learn Mem, 18(12), 742–746. doi:10.1101/lm.023671.111

Cohen, N. J., Poldrack, R. A., & Eichenbaum, H. (1997). Memory for items and memory for relations in the procedural/declarative memory framework. Memory, 5(1-2), 131–178.

Davachi, L. (2006). Item, context and relational episodic encoding in humans. Curr Opin Neurobiol, 16(6), 693–700. doi:S0959-4388(06)00152-8 [pii] 10.1016/j.conb.2006.10.012

Eacott, M. J., & Gaffan, E. A. (2005). The roles of perirhinal cortex, postrhinal cortex, and the fornix in memory for objects, contexts, and events in the rat. Q J Exp Psychol B, 58(3-4), 202–217. doi:TK0K3714113155J3 [pii] 10.1080/02724990444000203

Elfman, K. W., Parks, C. M., & Yonelinas, A. P. (2008). Testing a neurocomputational model of recollection, familiarity, and source recognition. J Exp Psychol Learn Mem Cogn, 34(4), 752–768. doi:10.1037/0278-7393.34.4.752

Guzowski, J. F., Knierim, J. J., & Moser, E. I. (2004). Ensemble dynamics of hippocampal regions CA3 and CA1. Neuron, 44(4), 581–584. doi:10.1016/j.neuron.2004.11.003

Hannula, D. E., Libby, L. A., Yonelinas, A. P., & Ranganath, C. (2013). Medial temporal lobe contributions to cued retrieval of items and contexts. Neuropsychologia. doi:10.1016/j.neuropsychologia.2013.02.011

Horner, A. J., Bisby, J. A., Bush, D., Lin, W. J., & Burgess, N. (2015). Evidence for holistic episodic recollection via hippocampal pattern completion. Nat Commun, 6, 7462. doi:10.1038/ncomms8462

Hsieh, L. T., Gruber, M. J., Jenkins, L. J., & Ranganath, C. (2014). Hippocampal activity patterns carry information about objects in temporal context. Neuron, 81(5), 1165–1178. doi:10.1016/j.neuron.2014.01.015

Huffman, D. J., & Stark, C. E. (2014). Multivariate pattern analysis of the human medial temporal lobe revealed representationally categorical cortex and representationally agnostic hippocampus. Hippocampus. doi:10.1002/hipo.22321

Johnson, M. K., Hashtroudi, S., & Lindsay, D. S. (1993). Source monitoring. Psychol Bull, 114(1), 3–28.

Knierim, J. J., Lee, I., & Hargreaves, E. L. (2006). Hippocampal place cells: parallel input streams, subregional processing, and implications for episodic memory. Hippocampus, 16(9), 755–764. doi:10.1002/hipo.20203

Kriegeskorte, N. (2011). Pattern-information analysis: from stimulus decoding to computational-model testing. Neuroimage, 56(2), 411–421. doi:S1053-8119(11)00097-8 [pii] 10.1016/j.neuroimage.2011.01.061

Kriegeskorte, N., Goebel, R., & Bandettini, P. (2006). Information-based functional brain mapping. Proc Natl Acad Sci U S A, 103(10), 3863–3868. doi:0600244103 [pii] 10.1073/pnas.0600244103

Kriegeskorte, N., Mur, M., & Bandettini, P. (2008). Representational similarity analysis - connecting the branches of systems neuroscience. Front Syst Neurosci, 2, 4. doi:10.3389/neuro.06.004.2008

Lacy, J. W., Yassa, M. A., Stark, S. M., Muftuler, L. T., & Stark, C. E. (2011). Distinct pattern separation related transfer functions in human CA3/dentate and CA1 revealed using high-resolution fMRI and variable mnemonic similarity. Learn Mem, 18(1), 15–18. doi:18/1/15 [pii] 10.1101/lm.1971111

LaRocque, K. F., Smith, M. E., Carr, V. A., Witthoft, N., Grill-Spector, K., & Wagner, A. D. (2013). Global similarity and pattern separation in the human medial temporal lobe predict subsequent memory. J Neurosci, 33(13), 5466–5474. doi:10.1523/JNEUROSCI.4293-12.2013

Leutgeb, J. K., Leutgeb, S., Moser, M. B., & Moser, E. I. (2007). Pattern separation in the dentate gyrus and CA3 of the hippocampus. Science, 315(5814), 961–966. doi:10.1126/science.1135801

Lever, C., Wills, T., Cacucci, F., Burgess, N., & O’Keefe, J. (2002). Long-term plasticity in hippocampal place-cell representation of environmental geometry. Nature, 416(6876), 90–94. doi:10.1038/416090a

Liang, J. C., Wagner, A. D., & Preston, A. R. (2012). Content Representation in the Human Medial Temporal Lobe. Cereb Cortex. doi:bhr379 [pii] 10.1093/cercor/bhr379

Libby, L. A., Hannula, D. E., & Ranganath, C. (2014). Medial Temporal Lobe Coding of Item and Spatial Information during Relational Binding in Working Memory. J Neurosci, 34(43), 14233–14242. doi:10.1523/JNEUROSCI.0655-14.2014

Maldjian, J. A., Laurienti, P. J., Kraft, R. A., & Burdette, J. H. (2003). An automated method for neuroanatomic and cytoarchitectonic atlas-based interrogation of fMRI data sets. Neuroimage, 19(3), 1233–1239.

Manns, J. R., & Eichenbaum, H. (2009). A cognitive map for object memory in the hippocampus. Learn Mem, 16(10), 616–624. doi:10.1101/lm.1484509

Marr, D. (1971). Simple memory: a theory for archicortex. Philos Trans R Soc Lond B Biol Sci, 262(841), 23–81.

McKenzie, S., Frank, A. J., Kinsky, N. R., Porter, B., Rivière, P. D., & Eichenbaum, H. (2014). Hippocampal representation of related and opposing memories develop within distinct, hierarchically organized neural schemas. Neuron, 83(1), 202–215. doi:10.1016/j.neuron.2014.05.019

Milivojevic, B., Vicente-Grabovetsky, A., & Doeller, C. F. (2015). Insight reconfigures hippocampal-prefrontal memories. Curr Biol, 25(7), 821–830. doi:10.1016/j.cub.2015.01.033

Mitchell, K. J., & Johnson, M. K. (2009). Source monitoring 15-years later: what have we learned from fMRI about the neural mechanisms of source memory? Psychol Bull, 135(4), 638–677. doi:10.1037/a0015849

Moore, M., Hu, Y., Woo, S., O’Hearn, D., Iordan, A. D., Dolcos, S., & Dolcos, F. (2014). A comprehensive protocol for manual segmentation of the medial temporal lobe structures. J Vis Exp(89). doi:10.3791/50991

Mumford, J. A., Davis, T., & Poldrack, R. A. (2014). The impact of study design on pattern estimation for single-trial multivariate pattern analysis. Neuroimage. doi:10.1016/j.neuroimage.2014.09.026

Mumford, J. A., Turner, B. O., Ashby, F. G., & Poldrack, R. A. (2012). Deconvolving BOLD activation in event-related designs for multivoxel pattern classification analyses. Neuroimage, 59(3), 2636–2643. doi:S1053-8119(11)01008-1 [pii] 10.1016/j.neuroimage.2011.08.076

Nichols, T. E., & Holmes, A. P. (2002). Nonparametric permutation tests for functional neuroimaging: a primer with examples. Hum Brain Mapp, 15(1), 1–25.

Norman, K. A. (2010). How hippocampus and cortex contribute to recognition memory: revisiting the complementary learning systems model. Hippocampus, 20(11), 1217–1227. doi:10.1002/hipo.20855

Norman, K. A., & O’Reilly, R. C. (2003). Modeling hippocampal and neocortical contributions to recognition memory: a complementary-learning-systems approach. Psychol Rev, 110(4), 611–646. doi:2003-08987-001 [pii] 10.1037/0033-295X.110.4.611

Poppenk, J., Evensmoen, H. R., Moscovitch, M., & Nadel, L. (2013). Long-axis specialization of the human hippocampus. Trends Cogn Sci, 17(5), 230–240. doi:10.1016/j.tics.2013.03.005

Ranganath, C. (2010). A unified framework for the functional organization of the medial temporal lobes and the phenomenology of episodic memory. Hippocampus, 20(11), 1263–1290. doi:10.1002/hipo.20852

Ranganath, C., & Ritchey, M. (2012). Two cortical systems for memory-guided behaviour. Nat Rev Neurosci, 13(10), 713–726. doi:10.1038/nrn3338

Ritchey, M., Libby, L. A., & Ranganath, C. (2015). Cortico-hippocampal systems involved in memory and cognition: the PMAT framework. Prog Brain Res, 219, 45–64. doi:10.1016/bs.pbr.2015.04.001

Ritchey, M., Montchal, M. E., Yonelinas, A. P., & Ranganath, C. (2015). Delay-dependent contributions of medial temporal lobe regions to episodic memory retrieval. Elife, 4. doi:10.7554/eLife.05025

Rolls, E. T., & Kesner, R. P. (2006). A computational theory of hippocampal function, and empirical tests of the theory. Prog Neurobiol, 79(1), 1–48. doi:10.1016/j.pneurobio.2006.04.005

Schlichting, M. L., Mumford, J. A., & Preston, A. R. (2015). Learning-related representational changes reveal dissociable integration and separation signatures in the hippocampus and prefrontal cortex. Nat Commun, 6, 8151. doi:10.1038/ncomms9151

Scoville, W. B., & Milner, B. (1957). Loss of recent memory after bilateral hippocampal lesions. J Neurol Neurosurg Psychiatry, 20(1), 11–21.

Vargha-Khadem, F., Gadian, D. G., Watkins, K. E., Connelly, A., Van Paesschen, W., & Mishkin, M. (1997). Differential effects of early hippocampal pathology on episodic and semantic memory. Science, 277(5324), 376–380.

Wills, T. J., Lever, C., Cacucci, F., Burgess, N., & O’Keefe, J. (2005). Attractor dynamics in the hippocampal representation of the local environment. Science, 308(5723), 873–876. doi:10.1126/science.1108905

Xue, G., Dong, Q., Chen, C., Lu, Z., Mumford, J. A., & Poldrack, R. A. (2010). Greater neural pattern similarity across repetitions is associated with better memory. Science, 330(6000), 97–101. doi:science.1193125 [pii] 10.1126/science.1193125

Yassa, M. A., & Stark, C. E. (2011). Pattern separation in the hippocampus. Trends Neurosci, 34(10), 515–525. doi:10.1016/j.tins.2011.06.006

